# Tissue structure accelerates evolution: premalignant sweeps precede neutral expansion

**DOI:** 10.1101/542019

**Authors:** Jeffrey West, Ryan O. Schenck, Chandler Gatenbee, Mark Robertson-Tessi, Alexander R. A. Anderson

## Abstract

Cancer has been hypothesized to be a caricature of the renewal process of the tissue of origin: arising from (and maintained by) small subpopulations capable of continuous growth^1^. The strong influence of the tissue structure has been convincingly demonstrated in intestinal cancers where adenomas grow by the fission of stem-cell-maintained glands influenced by early expression of abnormal cell mobility in cancer progenitors^2, 3^. So-called “born to be bad” tumors arise from progenitors which may already possess the necessary driver mutations for malignancy^4, 5^ and metastasis^6^. These tumors subsequently evolve neutrally, thereby maximizing intratumoral heterogeneity and increasing the probability of therapeutic resistance. These findings have been nuanced by the advent of multi-region sequencing, which uses spatial and temporal patterns of genetic variation among competing tumor cell populations to shed light on the mode of tumor evolution (neutral or Darwinian) and also the tempo^4, 7–11^. Using a classic, well-studied model of tumor evolution (a passenger-driver mutation model^12–16^) we systematically alter spatial constraints and cell mixing rates to show how tissue structure influences functional (driver) mutations and genetic heterogeneity over time. This model approach explores a key mechanism behind both inter-patient and intratumoral tumor heterogeneity: competition for space. Initial spatial constraints determine the emergent mode of evolution (Darwinian to neutral) without a change in cell-specific mutation rate or fitness effects. Driver acquisition during the Darwinian precancerous stage may be accelerated en route to neutral evolution by the combination of two factors: spatial constraints and limited cellular mixing.

---

Cancer is characterized by tens of thousands of somatic alterations, which may be classified into drivers (conferring advantageous, cancerous phenotypes to neoplastic cells) and passengers (neutral, nearly-neutral, or slightly deleterious mutations). Highly deleterious mutations are subject to negative selection and are removed from the population, while moderately deleterious mutations can evade purifying selection to remain present in an evolving tumor under selection pressure, a process known as “hitchhiking” with the sweeping driver clone^15, 17^.

Patterns of intratumoral genetic heterogeneity (ITH) and subclonal architecture are the direct consequence of the evolutionary dynamics of tumor growth. Multiregion sequencing has produced evidence that Darwinian evolution shapes at least part of ITH^10, 18^. Substantial increases in subclone fitness have been observed in some cases: 21% of colon cancers, 29% of gastric cancers, and 53% of metastases^19^. Early in tumorigenesis, selective sweeps are favored since early tumor growth must deal with obstacles such as spatial constraints, nutritional limitation, and immune attack^11^. However, the parental clone that can conquer these obstacles emerges from the subclones generated by early Darwinian evolution. This evolutionary bottleneck establishes the driver mutations in the parental, invasive clone as ubiquitous mutations in a neutral evolutionary phase of tumor growth, after invasion^11, 20^.

How does tissue structure influence somatic evolution? Some have speculated that modes of evolution (Darwinian to neutral) may be the outcome of cellular architecture of the tumor (e.g. the glandular structure of colorectal cancers limits the effects of selection), or of the malignancy’s anatomical location, governing access to resources or strong spatial constraints for growth^5, 19, 21^. Some structures are “amplifiers” of natural selection, improving the odds of advantageous mutants^22, 23^ (e.g. pancreatic ducts that control migration rates between patches^24^ and spatially segregated colon glands with a centralized stem cell pool^3^). The well-defined tissue structure begins to break down as the cancer transitions to increasing invasiveness, often with sudden, “punctuated” accrual of copy number alterations needed to facilitate invasion into the stroma^25, 26^. It is our point of view that this switch from Darwinian to neutral evolution is highly influenced by the tissue structure where the founding clone arises, and that transition to neutrality may occur early in tumor progression (before invasion).

The purpose of this manuscript is to model the effect of spatial constraints on the evolution of a precancerous tumor and apply this mathematical model to ductal carcinoma in situ (DCIS). We begin with a systematic and generalized understanding of the underlying mathematical model (in systematically varied spatial domain sizes and mixing rates) in figures 1 and 2. Next, this generalized understanding of the model is extended to a more biologically realistic setting: the 3-dimensional branching topology of a breast ductal network spatial structure (figure 3), representing biologically realistic timescales (500 to 1500 days), biologically realistic cell numbers for precancerous lesions (10^5^), and biologically realistic spatial topologies (ductal network structures).

**Figure 1.**
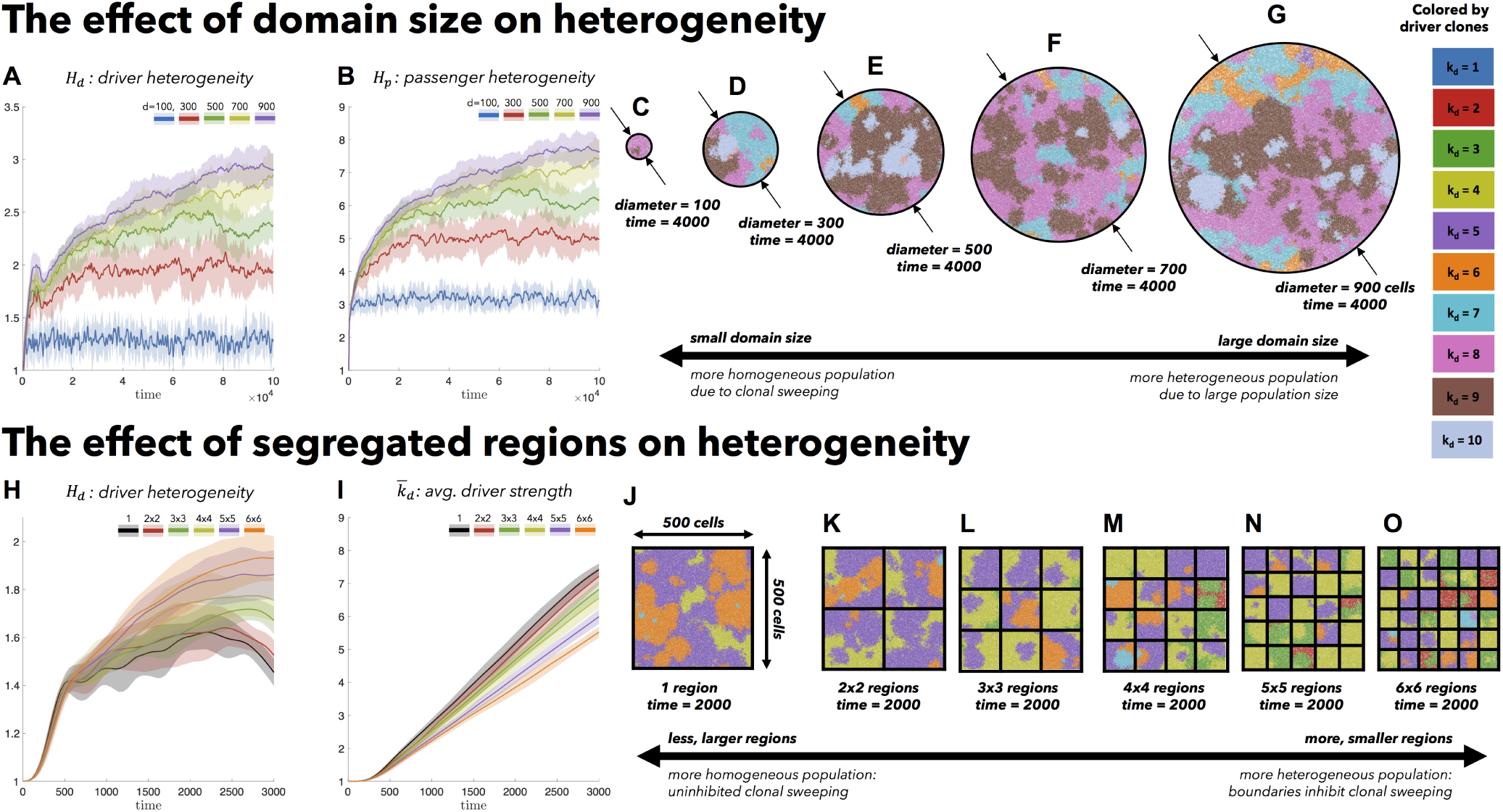
The effect of spatial constraints on heterogeneity: Cells divide and die on a regular square lattice. A cell selected for birth can divide only into an empty grid location and may accrue passenger or driver mutations. Top: simulations on varied sizes of domains, ranging from 100 cells in diameter to 900 cells, seeded with 100 cells (*k*_*d*_ = 1. *k*_*p*_ = 0) at time zero (*T*_*p*_ = 5 · 10^6^,*T*_*d*_ = 700,*s*_*d*_ = 0.1, *s*_*p*_ = 0.01). (A,B) An increase in domain size results in increased driver and passenger heterogeneity, with standard deviation shown (shaded colors) for 10 stochastic simulations. (C,D,E,F,G) Representative snapshots after 4000 cell generations. Bottom: Identical domain size (seeded with one-third of the domain filled; *k*_*d*_ = 1. *k*_*p*_ = 0) segregated into varied number of non-interacting regions (*T*_*p*_ = 10^6^, *T*_*d*_ = 700, *s*_*d*_ = 0.1, *s*_*p*_ = 0.01, *µ* = 10^−8^). (H) driver heterogeneity increases with number of segregated regions. (I) Boundaries limit new clones from expanding beyond a single region, decreasing the average number of drivers in the population. (J,K,L,M,N,O) Representative snapshots after 2000 cell generations. See attached video S3.

**Figure 2.**
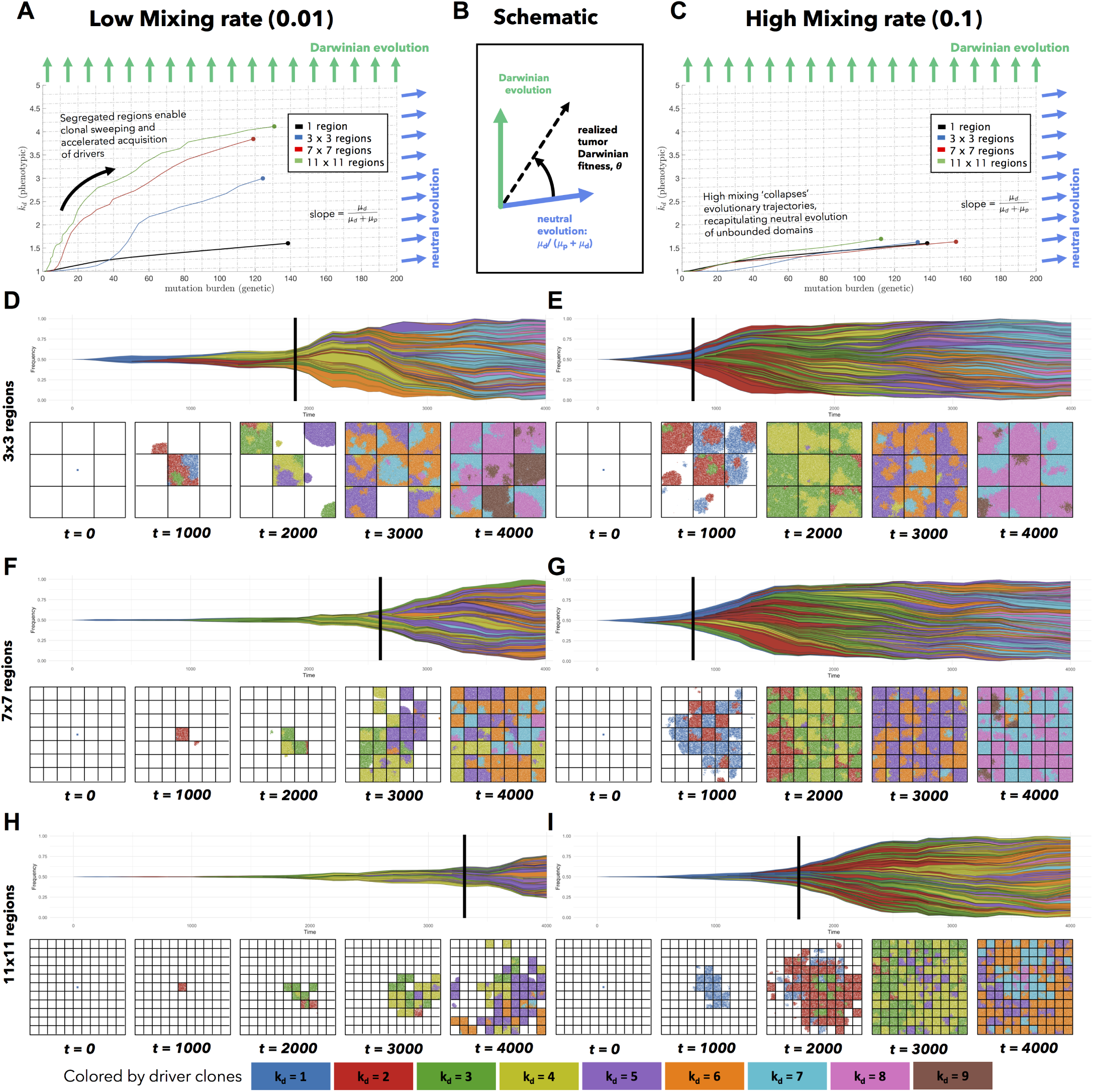
Spatial segregation with cell mixing accelerates evolution: (A,C) Tumors evolve on the genetic (mutation burden; x-axis) and phenotypic (average drivers; y-axis) axes. Simulations seeded with 100 cells (*k*_*d*_ = 1, *k*_*p*_ = 0) are allowed to mix between segregated regions at a low rate (0.01; left column) or high rate (0.1; right column) for varied number of segregated regions (line color). For low mixing, smaller regions impose higher selection pressure, accelerating the acquisition of drivers in the population (vertical axis). Simulations are run to identical tumor size (25% of the total domain). As mixing increases, tumor evolution “collapses” back onto the unsegregated single region case shown in black in B. (B) A tumor’s “realized fitness” can be quantified as the time-varying slope of the evolutionary trajectory in A and C. (C,D) A Muller plot of tumor evolution represents genotypes color-coded by driver (*k*_*d*_) value. The horizontal axis is time (cell generations), with height corresponding to genotype frequency. Descendant genotypes are shown emerging from inside their parents. Snapshots shown every 1000 cell generations. Simulations repeated for 7 by 7 regions (E,F), and 11 by 11 regions (G,H). Parameters: *T*_*p*_ = 5 · 10^6^, *T*_*d*_ = 700, *s*_*d*_ = 0.1, *s*_*p*_ = 10^−3^, *µ* = 10^−8^. See attached video S4.

**Figure 3.**
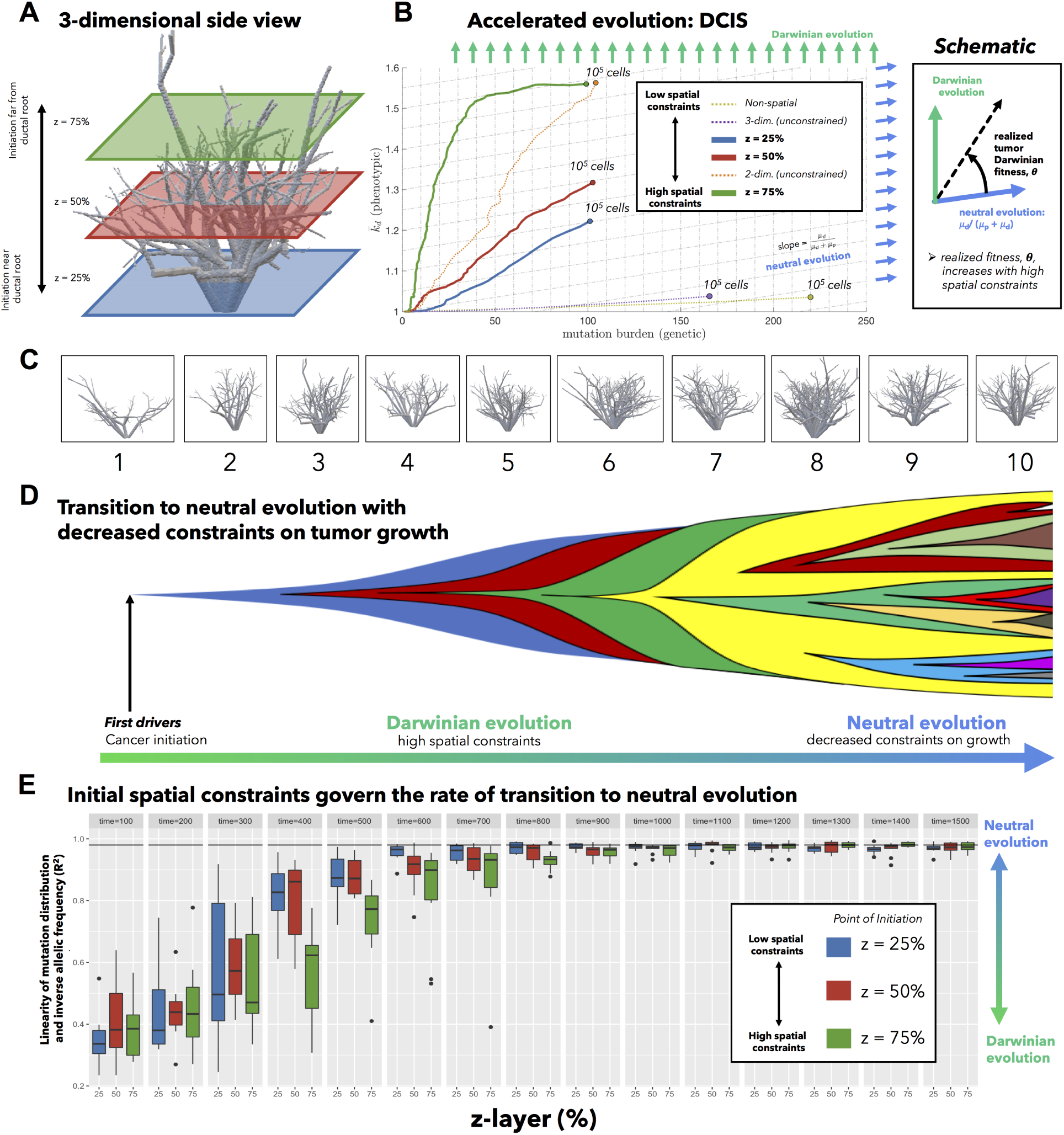
Three dimensional model of tumor evolution constrained by ductal network structure: (A) Realistic three-dimensional topology of breast ductal networks (reconstructed with data from anthropomorphic breast phantoms in^32^) provides full three-dimensional maps to seed and constrain tumor evolution simulations. (B) Tumor evolution is shown for varied points of initiation (z-dimension shown in A), to identical sizes (*N* = 10^5^ cells). See attached video S5. Each tumor is seeded with 500 cells (*k*_*d*_ = 1; *k*_*p*_ = 0, *T*_*p*_ = 10^6^, *T*_*d*_ = 700, *s*_*d*_ = 0.1, *s*_*p*_ = 0.1, *µ* = 10^−8^). The slope of trajectories (schematic) on this phase portrait is termed “realized tumor Darwinian fitness” and is dependent on spatial constraints at the point of tumor initiation. Simulations closer to the ductal root (e.g. blue curve, *z* = 25%) in larger, less constrained branches are characterized by a steady left-to-right (neutral) evolution and constant acquisition of new clones. Simulations further from the ductal root (e.g. green, *z* = 75%) in smaller, more constrained branches are characterized by clonal sweeping (bottom-to-top evolution) early, but with a shift toward neutrality (left-to-right) at later times. Trajectories may be compared to alternative models with identical parameterizations: non-spatial model (dashed yellow), unconstrained two-dimensional model (dashed orange), or unconstrained three-dimensional model (dashed purple). (C) The analysis is repeated subject to spatial constraints on ten highly distinct anthropomorphic breast phantom ductal network reconstructions. In general, ductal branches far from the root decrease in size and increase in number (see supplemental figure S7). (D) All simulations tend to follow an initially Darwinian evolutionary trajectory followed by a transition to neutral evolution. (E) An alternative metric of tumor neutrality: the linearity of cumulative mutation distribution with respect to inverse allelic frequency, sometimes called the 1/f power-law distribution^5^. Although all tumors initially are quantitatively shown to be non-neutral, all eventually progress to neutral or nearly-neutral state (the neutral minimum threshold of 0.98^5^ shown in black). Tumors with initially high spatial constraints (green bars) transition to neutral evolution more slowly than those with less spatial constraints (blue bars). High spatial constraints enable accelerated evolution over a range of ductal networks in C (see supplemental figure S9).

## Functional and genetic heterogeneity

While many mathematical passenger-driver tumor evolution models use branching processes^27^ or stochastic, well-mixed (non-spatial), agent-based models^12, 13, 21, 28, 29^, some have investigated the role of *spatial* competition on local heterogeneity and circulating tumor cells^16^, resistance to therapy^30^, metastasis^6, 14^, and trade-offs between migration and proliferation^31^. The model introduced here is a spatially-explicit extension of a previously published non-spatial model of passengers and drivers in tumor evolution^12, 13, 15^. Tumor evolution is played out on a two- or three-dimensional grid where each grid point can contain at most one cell. Each cell carries heritable genetic changes classified into driver mutations (e.g. an activating mutation in KRAS) or passenger mutations. Cells begin each simulation with a single driver mutation, where each subsequent driver increases the birth rate (i.e. multiplicative epistasis) by a factor of a fitness advantage parameter, *s*_*d*_. Similarly, passenger mutations decrease the birth rate by a factor of fitness penalty, *s*_*p*_ (see figures S1 and S2). We begin by demonstrating some results of the model in varied spatial domain sizes and cellular mixing rates in 2-dimensions (see figures 1 and 2) and subsequently apply these conceptual findings to ductal carcinoma in situ (figure 3).

Tumors constrained to smaller domain sizes (figure 1, top; video S3, top) show consistently lower driver and passenger diversity than for larger domain sizes. Small, tightly-coupled homogeneous populations of cells are able to quickly sweep each successive driver mutation. Larger domains consist of a heterogeneous population; with many more cell divisions, the odds of accruing another driver mutation are increased, but they have little chance of sweeping through the large domain. While differences in heterogeneity measures (figure 1A,B) diverge quickly for varied domain sizes, they do not approach steady states until extreme time scales (20,000 generations; 50 years). This provides a baseline comparison for simulations in biologically realistic spatial settings (and biologically realistic timescales), shown below.

To control for population size effects in this evolutionary arms race, identical domains are segregated into non-interacting regions of varying size (figure 1, bottom). Again, smaller (highly constrained) regions provide a stronger selection force, sweeping away weaker subclones. This sweeping stops at each region’s boundaries, resulting in a heterogeneous population of locally homogeneous regions (figure 1H). Boundaries limit new clones from expanding beyond a single region, decreasing the average number of drivers in the population (figure 1I).

## Modeling the acceleration of evolution by tissue structure

Bounded, non-interacting regions play a role in human cancers, which are often locally constrained to a single gland or a duct. Such glandular or ductal structures allow for limited cellular mixing during premalignant growth, enabling the tumor to explore new (and often less constrained) environments. In figure 2, each segregated region may now circulate cells into a neighboring region at a low or high rate of mixing (left and right columns, respectively). Tumors evolve on two axes: genetic (mutation burden; x-axis) and phenotypic (average number of driver mutations,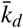; y-axis). The evolution of an unsegregated tumor in is shown in black (figure 2A,C), accumulating genetic diversity (left-to-right) over time with a slow accumulation of drivers (bottom-to-top).

This state space diagram allows us to track the accelerated acquisition of drivers in a precancerous population, with respect to tumor size. The trajectory of a tumor’s evolution quantifies the tumor-scale effect of domain size and mixing, over time. Despite seeding simulations with identical parameterization, tumors may evolve in a neutral or Darwinian mode (or on a continuous scale between the two, shown in figure 2B), subject to selection imposed by domain and cellular mixing. Neutral tumors acquire drivers at a rate equal to the ratio of drivers to all mutations (figure 2A,C, blue arrows). Conversely, Darwinian tumors sweep each new driver mutation through the population, resulting in a vertical trajectory (figure 2A,C green arrows).

Here we propose a novel classification of time-dependent tumor evolution as its “realized Darwinian fitness,” the time-varying angle on the mutation-driver state space (see figure 2B). Tumors are able to realize much higher fitness levels (figure 2A) with high spatial constraints and limited mixing, enabling rapid acquisition of drivers, despite no changes in subclonal fitness effects.

This view of Darwinian fitness represents a paradigm shift in tumor neutrality. Previous work has often focused on inferring cell-specific fitness (i.e. subclonal selection) through variant allele frequency distribution metrics (e.g. the 1/f power law distribution)^5, 33, 34^. Alternatively, we argue that there is a new scale of fitness that should be considered: tumor-scale. We view this as a paradigm shifting insight: the surrounding spatial context modulates the “realized tumor fitness.” Spatial constraints accelerate evolution, without changes in cell-specific fitness.

Tumor diversity can be visualized on a Muller plot, displaying each clone’s abundance over time, colored by driver mutations (figure 2D - I; video S4). Tumors with low mixing rates (figure 2, left column) tend to evolve by a Darwinian mode of evolution: clonal sweeping which maintains a lower genetic diversity. Increasingly smaller regions allows new drivers to more easily sweep within a local region (figure 2A). Introducing segregated regions increases spatial constraints, allowing the tumor to realize increased levels of selection (figure 2, left column) even without changes in cell-specific fitness. Importantly, this accelerated mode of evolution is lost when mixing between regions is too high (see figure 2C). The high mixing rate (right column) recapitulates the evolutionary trajectory of a relatively unconstrained tumor, decreasing the tumor’s realized fitness.

## Ductal Carcinoma in situ

The previous two figures provide a systematic understanding of the model behavior, so we now we turn our attention to an application of the importance of spatial constraints in premalignant evolution. The branching topology of a breast ductal and glandular network structure acts as a evolutionary accelerant, where spatially segregated regions (ductal branches) work in combination with cell mixing (subject to varied branching topology) to accelerate tumor evolution^35^. In figure 3, the evolution of ductal carcinomas in situ is initialized and constrained to grow inside a ductal network reconstructed with data from anthropomorphic breast phantoms^32^. This provides a realistic three-dimensional topology of a continuously connected series of progressively smaller branches (figure 3A; video S6).

Tumors initiated inside larger ductal branches near the root of the network (*z* = 25%, blue, figure 3B) begin with less spatial constraints and increased access to expand into new branches. These tumors evolve more neutrally (left-to-right trajectories in figure 3B). Tumors initiated further from the ductal root in smaller, more constrained branches (e.g. purple curve) are characterized by clonal sweeping (vertical trajectories) early. At later times, the tumor expands into new unexplored territories, shifting toward neutral trajectories. A tumor originating in a tightly constrained duct enables an accelerated acquisition of drivers early in tumor progression (10^5^ cells), which may be a more dangerous, highly homogeneous population of malignant cells that have all acquired new traits such as invasiveness, motility, or metastatic capabilities. These important conclusions are lost when considering identical parameterizations of a non-spatial model (figure 3B, yellow), unconstrained two-dimensional model (orange), or unconstrained three-dimensional model (purple).

Surprisingly, two otherwise identical tumors may realize dramatic differences in fitness depending on constraints imposed by tissue architecture. On the cell scale, any given subclone may indeed have a selective advantage (i.e. a higher birth rate). Yet, the effective outcome of this subclonal advantage depends on the surrounding competitive context of that cell. In other words, cell-specific phenotypic behavior can be “overridden” by the tissue architecture, allowing the tumor to realize increased fitness. This approach adds clarity to the debate of neutral tumor evolution by exploring a key mechanism behind both inter-patient and intratumoral tumor heterogeneity: competition for space.

Neutral evolution has previously been quantified using the distribution of the cumulative number of mutations in the tumor with respect to inverse allelic frequency (1/f power-law distribution)^5^. We simulate sequencing (over time) and quantify the linearity of the 1/f power-law distribution (figure 3E) and repeat the analysis subject to spatial constraints on ten anthropomorphic breast phantom ductal network reconstructions shown in figure 3C. Although all tumors (regardless of spatial constraints) initially are quantitatively shown to be non-neutral (figure 3E), all eventually progress to a neutral or nearly-neutral evolutionary mode (the neutral minimum threshold of 0.98^5^ shown in black). Initial high spatial constraints (green bars) transition to neutral evolution more slowly than lack of spatial constraints (blue). En route to neutral evolution, high spatial constraints enable accelerated evolution by two processes (see supplemental figure S9). First, mutation burden is limited via negative selection. Second, tumor growth is limited, enabling easier subclonal expansion of more fit clones into a smaller population.

It is clear that spatial competition can no longer be ignored in evolutionary models of tumor evolution. These results (summarized in figure 4) help to unify the debate surrounding neutral tumor evolution by clarifying the role of space in the transition from Darwinian to neutral evolution. Initial spatial constraints determine the emergent mode of evolution (neutral to Darwinian) without the necessity for changes in cell-specific mutation rate or fitness effects. The branching topology of ductal networks at tumor initiation determines two important evolutionary accelerants: spatial constraints and cellular mixing. This connectivity is likely to be highly heterogeneous between patients, leading to variability in rates of cellular mixing between spatially distinct niches within a tumor. Although all tumors tend toward neutrality, spatial constraints allow tumors to linger in the non-neutral mode for longer, acquiring more drivers per cell. Limited connectivity enables subclones to undergo high levels of local selection due to spatial constraints. Our new metric of realized fitness emphasizes the need to consider both space and time when inferring the mode of evolution. Whilst this type of quantification is not currently possible for clinical tissues, our results indicate that we must be cautious when interpreting non-spatial measures of evolution.

**Figure 4.**
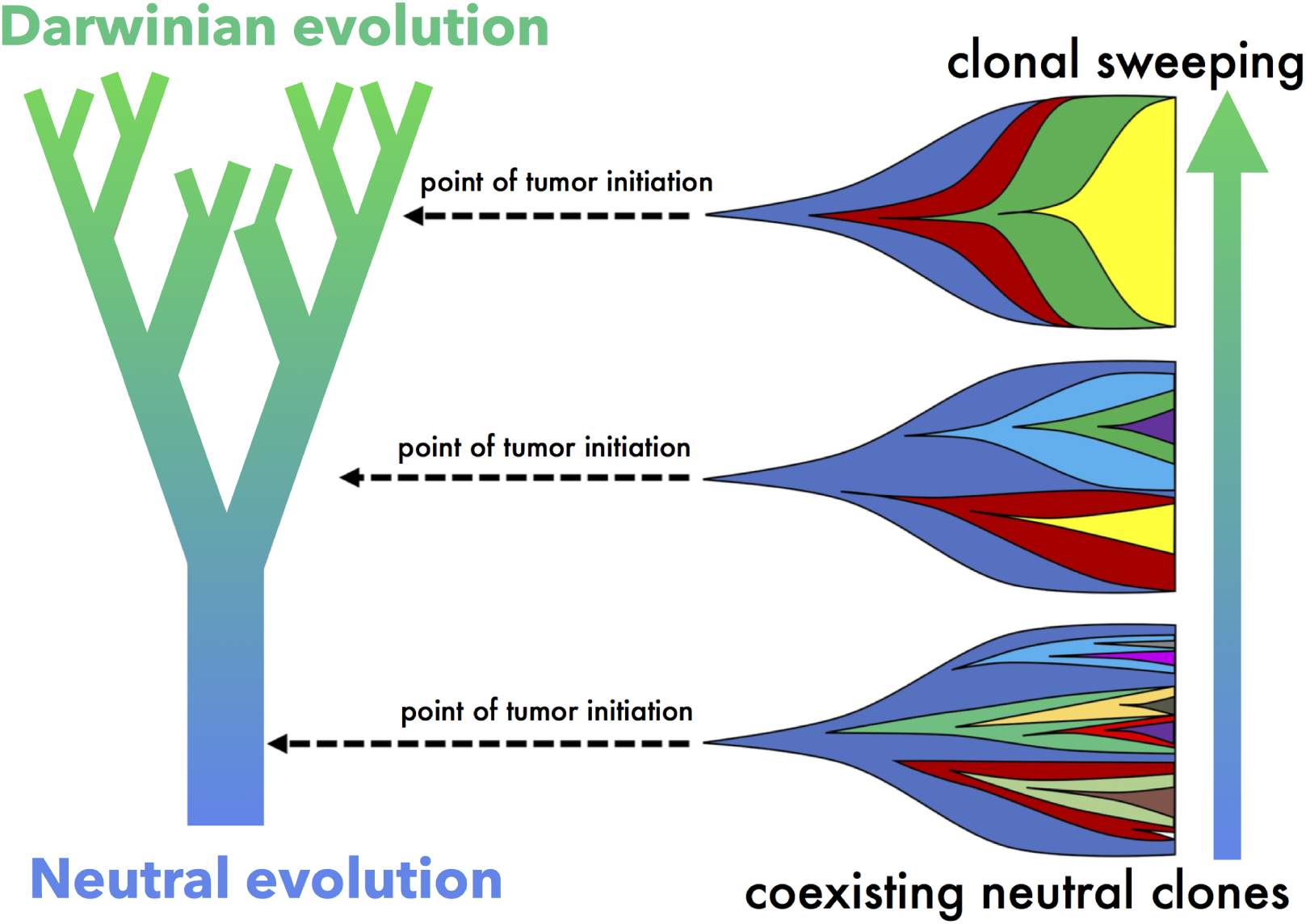
Summary schematic: Simulations near the ductal root in larger, less constrained branches (e.g. bottom) are characterized by a steady, neutral evolution, consistently acquiring new clones that coexist for extended time periods. A smaller tumor originating in a smaller, more constrained branches far from the ductal root (e.g. top) enables an accelerated evolution with clonal sweeping. Levels of selection may be highly heterogeneous between patients, due to the location of tumor initiation.

## Methods

Consistent with previously models of passenger driver evolution, tumors will undergo progression with a low mutation rate (less total deleterious passengers) or a low passenger fitness *s*_*p*_ (see figure S1A,B). We specifically are focused on measures of functional (drivers) and non-functional (passengers) heterogeneity and the conclusions drawn from the model apply in the assumption of neutral or non-neutral passengers.

### Code Availability

All figures were produced using an agent-based modeling platform (Java) known as HAL: Hybrid Automata Library^36^ (available at: https://github.com/MathOnco/HAL).

### Model Overview

Each model simulation is carried out on a two- or three-dimensional grid lattice where each tumor cell is allowed to occupy a single grid point. Simulations are started with 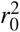 initial cells (*r*_0_ = 10 unless otherwise noted). During each time step, each cell undergoes a birth-death process with the following birth (*P*_*b*_) and death (*P*_*d*_) probabilities:

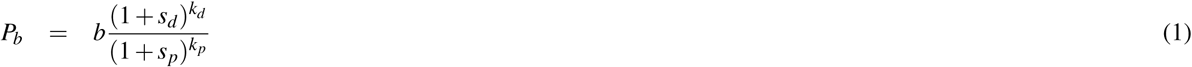

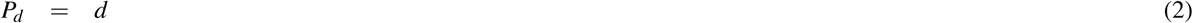

where *b* and *d* are the baseline birth and death rates, respectively. Tumor cells are initiated with exactly one driver mutation (i.e. *k*_*d*_ = 1) and zero passenger mutations (i.e. *k*_*p*_ = 0). During the birth process, cells may undergo mutations at a rate *µ*_*d*_ = *T*_*d*_*µ* (driver mutations) and *µ*_*p*_ = *T*_*p*_*µ* (passenger mutations). The model is a spatially-explicit extension of ref.^12^ where a driver mutation is a rare event with confers a fitness advantage to birth rate known as *s*_*d*_ and a passenger mutation is relatively common event that confers some fitness penalty, *s*_*p*_. Here we make the simplifying assumption that each subsequent driver (and each subsequent passenger) has equal effect, rather than a distribution of fitness effects, but others have shown that relaxing this assumption gives similar dynamics^12^.

### Model parameters

Consistent with the range reported^12, 13^, the following parameter values were used: *s*_*d*_ = 0.1, *s*_*p*_ = 10^−3^, *T*_*d*_ = 700, *T*_*p*_ = 10^6^, *µ* = 10^−8^. Due to the possibility of extensive variability in these parameters (see ref.^13^), each of these parameters was varied several orders of magnitude to ensure the robustness of conclusions drawn. Birth and death rates were kept constant at *b* = *d* = 0.5 unless otherwise noted, and do not significantly alter results for *b, d ∈* [0.1, 0.5] (see figure S1B).

### Heterogeneity

Heterogeneity of driver and passenger mutations is calculated using Shannon entropy, given by:

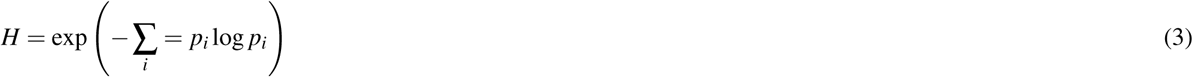

where *p*_*i*_ is the proportion of cells within the population with exactly *i* driver (*H*_*d*_) or passenger (*H*_*p*_) mutations.

### Dispersal rate between glands and ducts

Because there are no reliable data on the probability of tumor cell mixing and dispersal between breast ducts, a range of dispersal rates (rate ∈ [0.01,0.1]) are simulated in dimensions, while realistic breast ductal network structure in three dimensions with varied ductal branch sizes and branching topology (with varied initial conditions) are given by ref.^32^.

### Quantification of ductal number and area

Simulations were performed subject to spatial constraints on ten anthropomorphic breast phantom ductal network reconstructions, shown in figure S7, below. For each two-dimensional slice (e.g. figure S7B), the number of ductal branches are counted and quantified using OpenCV and scikit-image for the Python programming language. In order to minimize bias introduced when ductal branches run parallel to a given slice, an ellipse is fit (figure S7B, bottom) to each ductal branch to find the length of the minimum axis, *d*_min._. Distributions of *d*_min._ are shown in figure S7A for each range of z-values (colored purple, gray, pink, yellow) and repeated for ten breast ductal network structures. In general, there are fewer, larger ducts near the root of the ductal network and many, smaller ducts as z-layer increases (figure S7C).

## Supplementary Figures

**Figure S1.**
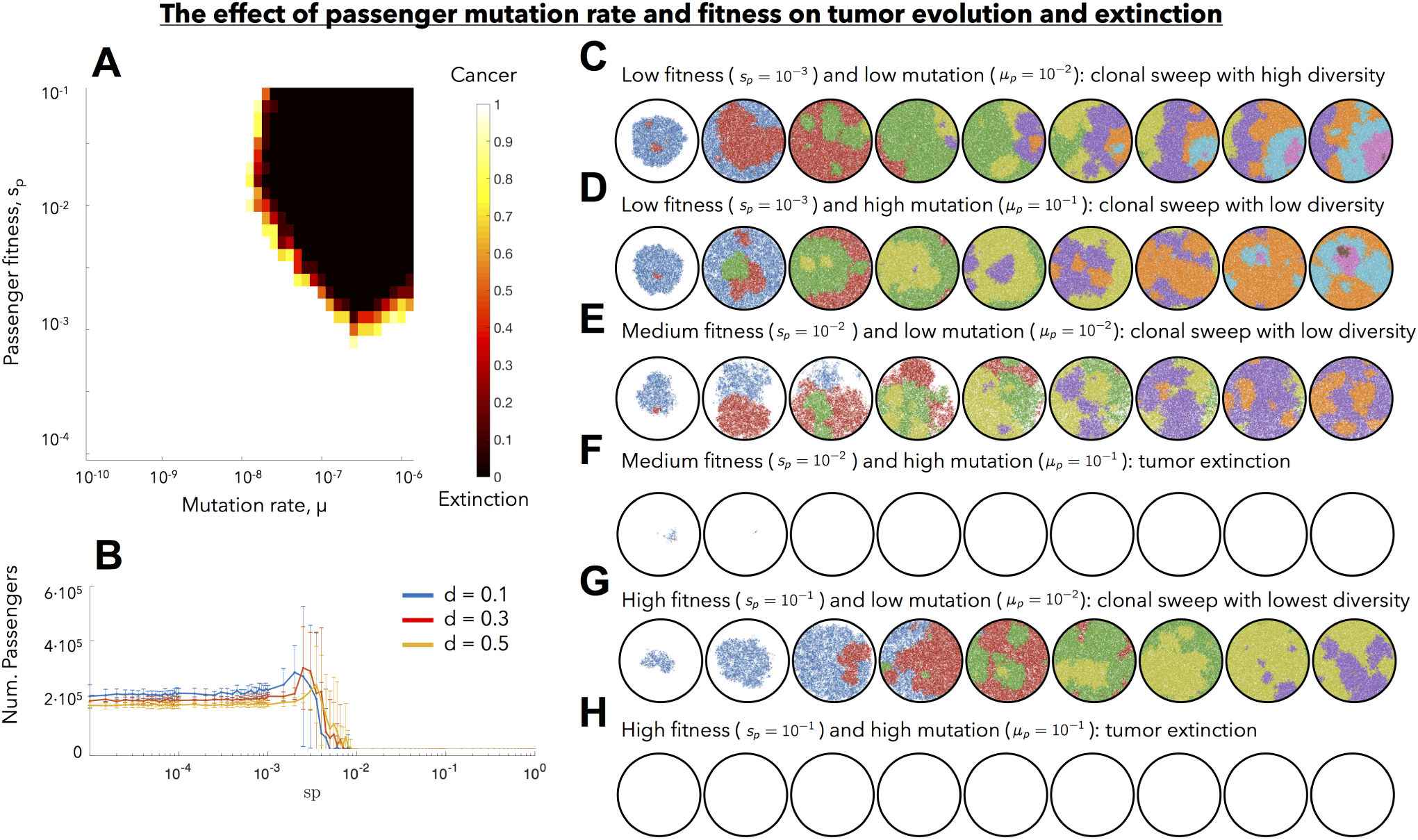
Cancer progression and extinction in passenger driver evolution. (A) Rate of cancer progression (simulations that do not go extinct after 10000 cell generations) and extinction depends on passenger fitness penalty, *s*_*p*_ and mutation rate, *µ*. 10 simulations shown for each parameter value. (B) Low and moderately deleterious passenger mutations can evade negative selection, for a variety of turnover (cell death, *d*) values. (C) Low passenger fitness penalty and low mutation rate leads to clonal sweep with high diversity at each time point. (D) Low passenger fitness and high mutation rate leads to clonal sweep with lower diversity. (E) Moderate passenger fitness and low mutation rate similarly leads to clonal sweep with lower diversity. (F) Moderate passenger fitness with high mutation rate leads to extinction. (G) High passenger fitness with low mutation rate leads to clonal sweep with lowest diversity. (H) High passenger fitness with high mutation rate leads to tumor extinction.

**Figure S2.**
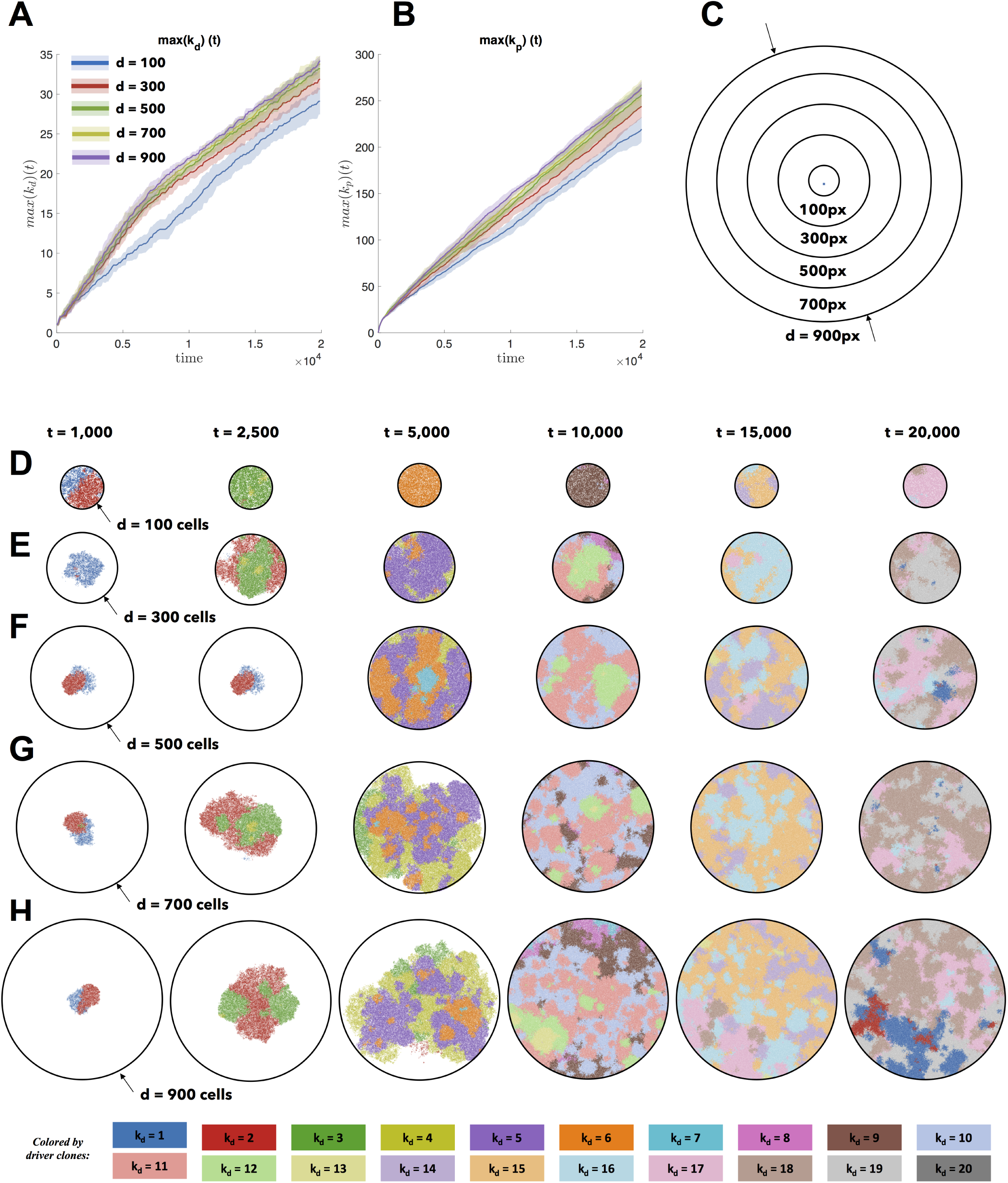
The effect of spatial competition on passenger hitch-hiking. The maximum driver number in the population, max(*k*_*d*_) and the maximum passenger, max(*k*_*p*_) is shown for a range of size constraints. Apart from the smallest domain size (D) of 100 cells, the maximum driver and passenger for each size constraint are relatively equivalent for each point in time. (C) Domain sizes, shown to scale. (D,E,F,G,H) As seen from the snapshots at each time point the population ranges from heterogeneous (large domains) to homogeneous (smaller domains).

## Supplemental Video Captions

**S3 (S3.mov) The effect of spatial constraints on heterogeneity (video)** Identical simulations and parameterization as figure 1. Cells divide and die on a regular square lattice. A cell selected for birth can divide only into an empty grid location and may accrue passenger or driver mutations. Top: simulations on varied sizes of domains, ranging from 100 cells in diameter to 900 cells, seeded with 100 cells at time zero (*k*_*d*_ = 1. *k*_*p*_ = 0) at time zero (*T*_*p*_ = 5 · 10^6^,*T*_*d*_ = 700,*s*_*d*_ = 0.1, *s*_*p*_ = 0.01). Bottom: Identical domain size (seeded with one-third of the domain filled; *k*_*d*_ = 1. *k*_*p*_ = 0) segregated into varied number of non-interacting regions (*T*_*p*_ = 10^6^, *T*_*d*_ = 700, *s*_*d*_ = 0.1, *s*_*p*_ = 0.01, *µ* = 10^−8^).

**S4 (S4.mov) Spatial segregation with cell dispersal accelerates evolution (video)** Identical simulations and parameterization as figure 2. Simulations seeded with 100 cells (*k*_*d*_ = 1, *k*_*p*_ = 0) are allowed to disperse between segregated regions at a low rate (0.01; left column) or high rate (0.1; right column) for varied number of segregated regions, as shown. A Muller plot of tumor evolution represents genotypes color-coded by driver (*k*_*d*_) value. The horizontal axis is time (cell generations), with height corresponding to genotype frequency. Descendant genotypes are shown emerging from inside their parents. Simulations repeated for 7 by 7 regions, and 11 by 11 regions. Parameters: *T*_*p*_ = 5 · 10^6^, *T*_*d*_ = 700, *s*_*d*_ = 0.1, *s*_*p*_ = 10^−3^, *µ* = 10^−8^.

**S5 (S5.mov) Three dimensional model of tumor evolution constrained by ductal network structure (video)** Identical simulations and parameterization as figure 3. Realistic three-dimensional topology of breast ductal networks (reconstructed with data from anthropomorphic breast phantoms in^32^) provides full three-dimensional maps to seed and constrain tumor evolution simulations. Tumor evolution is shown for varied initial conditions (*T*_*p*_ = 10^6^, *T*_*d*_ = 700, *s*_*d*_ = 0.1, *s*_*p*_ = 0.1, *µ* = 10^−8^). Simulations closer to the ductal root (top) in larger, less constrained branches are characterized by more consistently neutral evolution and constant acquisition of new clones. Simulations further from the ductal root (bottom) in smaller, more constrained branches are characterized by clonal sweeping early, but neutral evolution at later times. Each tumor is seeded with 500 cells (*k*_*d*_ = 1; *k*_*p*_ = 0). Each simulation is run for 3000 cell generations.

**S6 (S6.mov) Realistic three-dimensional topology of breast ductal networks (video)** Left: realistic three-dimensional topology of breast ductal networks, reconstructed with data from anthropomorphic breast phantoms in^32^ provides full three-dimensional maps to seed and constrain tumor evolution simulations. This provides a topology of a continuously connected series of progressively smaller ductal branches. Middle: slices in the z-dimension show fewer, larger ducts for low z-values and many, smaller ducts for high z-values. Right: static images of z-dimension slices.

**Figure S7.**
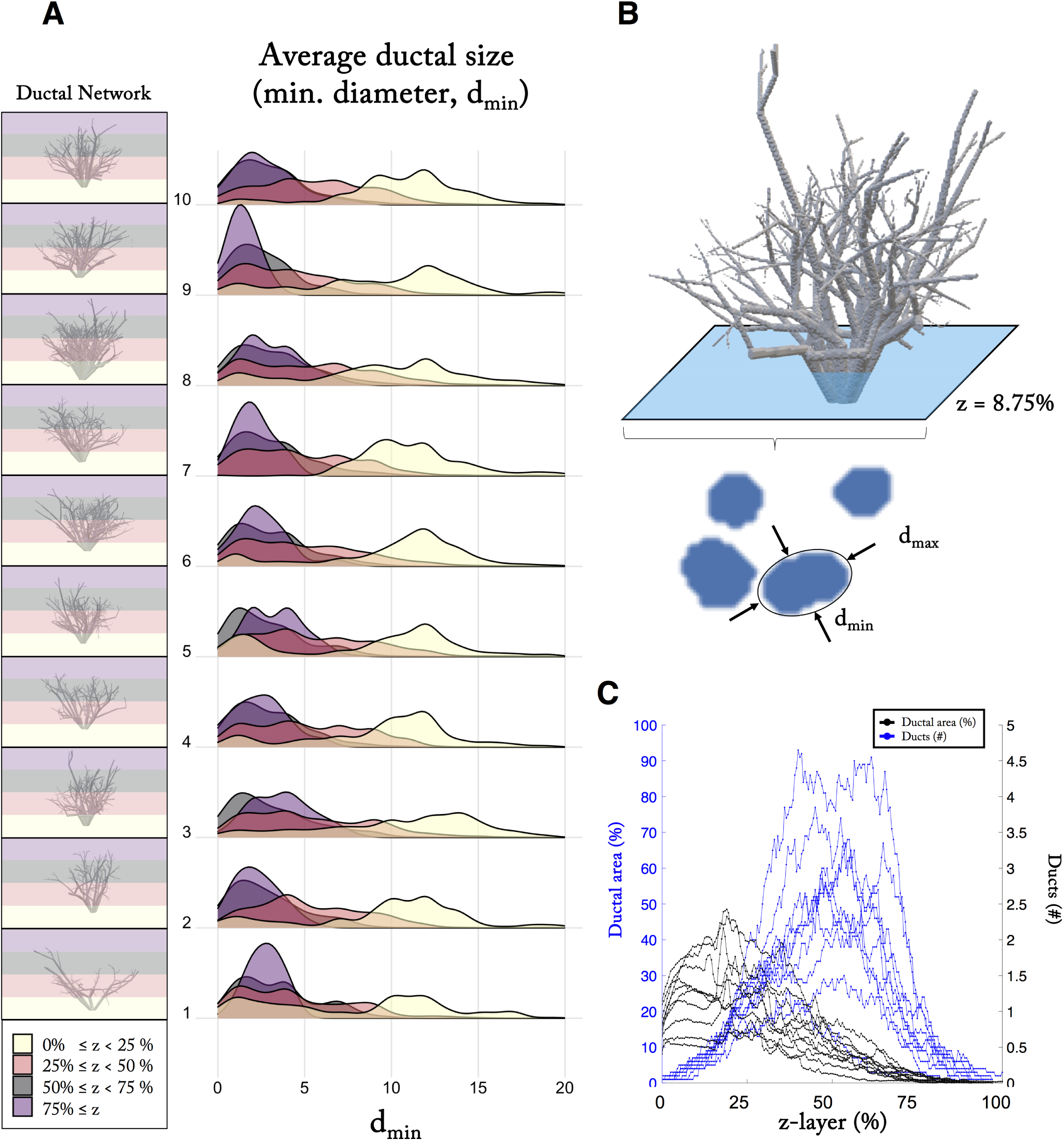
Quantification of spatial constraints imposed by breast ductal networks. (A) Data from ten anthropomorphic breast phantom ductal network reconstructions. Distributions of average minimum diameter (shown in B) are shown for varied z (relative distance from ductal root). (B) For each two-dimensional slice, the number of ductal branches are counted and quantified using Python OpenCV. In order to minimize bias introduced when ductal branches run parallel to a given slice, an ellipse is fit (figure S8B, bottom) to each ductal branch. (C) Ductal area (percentage) and number of ducts is shown as a function of z (relative distance from ductal root). In general, there are fewer, larger ducts near the root of the ductal network and many, smaller ducts as z-layer increases.

**Figure S8.**
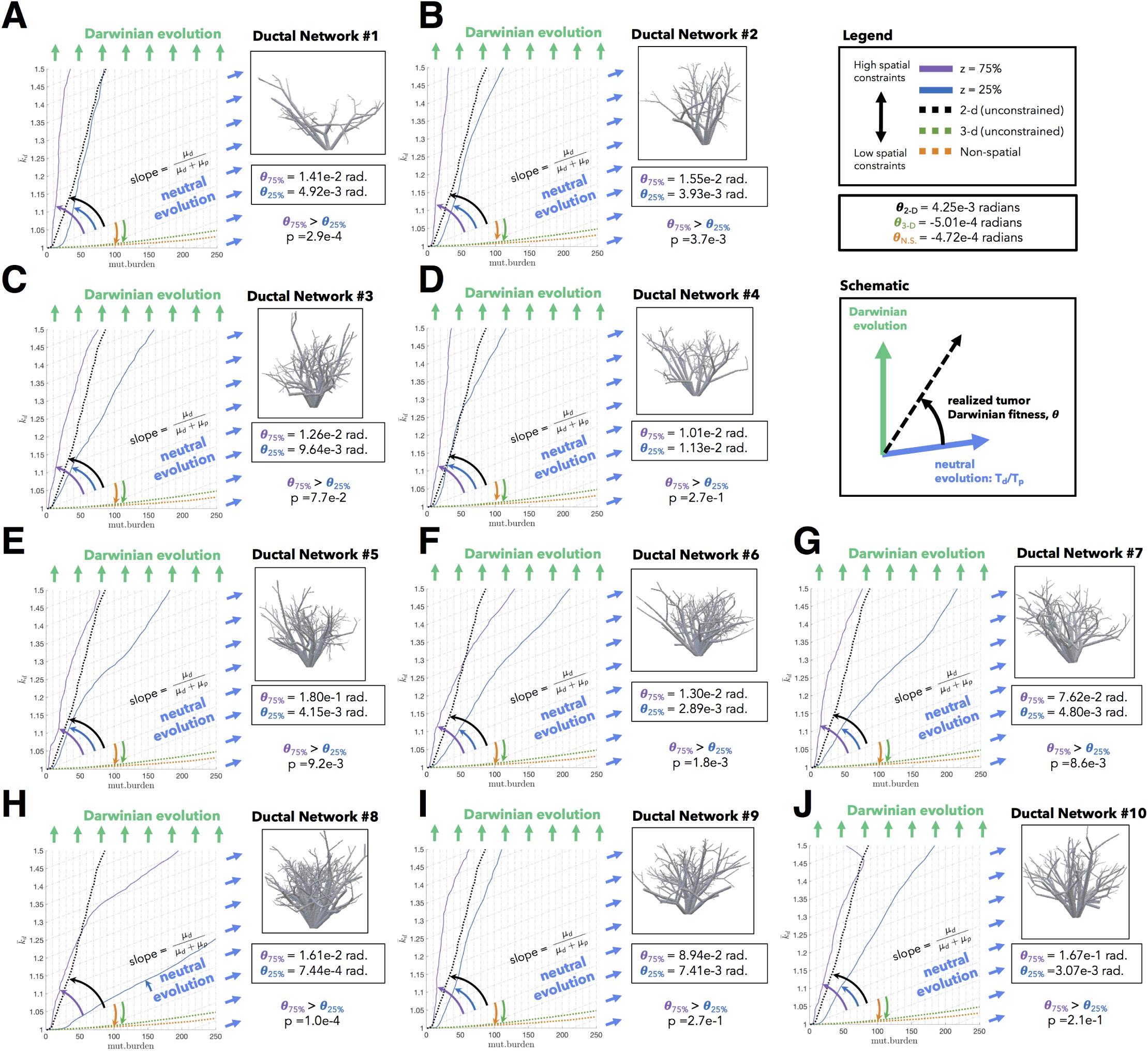
Statistical significance of realized tumor fitness, *θ* with varied initial spatial constraints. Data from ten anthropomorphic breast phantom ductal network reconstructions (shown in S7) constrain tumor growth in simulations. Trajectories are shown initiated in high spatially constrained regions of the ductal network (*z* = 75%) and low spatially constrained regions (*z* = 25%). Angles and p-values (one-tailed t-test) are reported in (A) through (J). These important conclusions are lost when considering identical parameterizations of a non-spatial (orange dashed line), two-(black dashed line), or three-dimensional (green dashed line) model.

## Acknowledgments

The authors gratefully acknowledge funding from the Physical Sciences Oncology Network (PSON) at the National Cancer Institute, U54CA193489 (supporting JW, CG, MRT, ARAA). ROS is supported by the Wellcome Trust (grant no. 108861/7/15/7) and the Wellcome Centre for Human Genetics (grant no. 203141/7/16/7). The authors gratefully acknowledge Hanbean Youn for helpful feedback and communication, as well as providing access to data from anthropomorphic breast phantoms used to reconstruct breast ductal networks^32^.

## Author Contributions

JW and ARAA conceived the research question and model design. JW performed analysis of data from the model and wrote the manuscript with critical comments and input from ARAA and MRT. CG and ROS developed code used to visualize Muller plots and assisted with manuscript edits. All authors have read and edited the final manuscript.

